# The tardigrade *Hypsibius exemplaris* as an emerging model to study the mitochondrial alternative oxidase at animal organismal level

**DOI:** 10.1101/2020.12.09.417485

**Authors:** Daria Grobys, Milena Roszkowska, Łukasz Kaczmarek, Wiesława Jarmuszkiewicz, Andonis Karachitos, Hanna Kmita

## Abstract

Mitochondrial alternative oxidase (AOX) is present in mitochondria of many invertebrates. Independently of the reason concerning the enzyme occurrence in animal mitochondria, expression of AOX in human mitochondria is regarded as a potential therapeutic strategy. Till now, relevant data were obtained due to heterologous AOX expression in cells and animals without natively expressed AOX. Application of animals natively expressing AOX importantly contribute the research. Thus, we investigated *Hypsibius exemplaris* as a model for AOX activity analysis. We observed that *H. exemplaris* tolerance to the blockage of the MRC complexes was diminished in the presence of AOX inhibitor and the inhibitor-sensitive respiration enabled the tardigrade respiration under condition of the blockage. Furthermore, although detection of AOX at protein level and pronounced oxygraphic registration of its activity required the MRC complex blockage, the obtained data indicated that AOX clearly contributed animal functioning. We demonstrated that AOX activity in tardigrades, can be monitored by measurement of intact specimen whole-body respiration. Furthermore, it was also possible to monitor the impact of the MRC complex IV blockage on AOX expression and AOX inhibition in the absence of the blockage on animal functioning. Thus, *H. exemplaris* is applied as a whole-animal model suitable to study AOX.

## Introduction

Currently, it is suggested that the mitochondrial respiratory chain (MRC) of most invertebrates contains the alternative oxidase (AOX), an enzyme which provides a secondary oxidative pathway to the classical cytochrome pathway (e.g. [1]) but till now, natively expressed AOX level and activity have not been detected simultaneously in intact animals. Since the enzyme is not present in mitochondria of vertebrates, it is hypothesized that AOX-based respiration was critical to the evolution of animals by enabling oxidative metabolism during the transition to a fully oxygenated Earth [2]. Thus, AOX may be treated as an adaptation to oxygen shortage which, in present-day organisms, may be caused by MRC deficiency or blockage. This in turn implies AOX application in treatment of mitochondrial diseases. As summarized by Saari et al. [3], the diseases may range from primary mitochondriopathies to common disease entities where mitochondrial disruption is due to ischemia/reperfusion injury, oxidative or proteotoxic stress, toxic damage or other external causes. However, the relevant data has been obtained by heterologous AOX expression in cells and animals that do not have native AOX although it appears to be essential for understanding of AOX contribution to animal physiology and consecutive development of AOX-based therapeutic strategies. The AOX activity can be understand by detailed investigation of effects imposed by heterologous AOX expression on physiological responses of model organisms under stressful environmental conditions [3]. However, the research could be greatly simplified by application (as a model) of animals natively expressing AOX.

According to the definition, AOX is the mitochondrial inner membrane enzyme introducing a branch into the canonical animal MRC formed by four main multi-subunit complexes numbered from I to IV (Fig. 1). The branching occurs before the MRC complex III, at the ubiquinone/ubiquinol pool, and results in transferring of electrons to oxygen with sustained proton pumping by the MRC complex I but without proton pumping by the MRC complexes III and IV. The transfer is antimycin A (AA)- and cyanide-insensitive because AOX is not inhibited by AA and cyanides which are frequently used as inhibitors of the MRC complexes III and IV, i.e. ubiquinol–cytochrome c reductase and cytochrome c oxidase, respectively (e.g. [1, 4-8]). Consequently, proton-pumping by MRC is confined to complex I which results in cyanide- and AA-resistant respiration. Since proton gradient, formed due to proton pumping, is required for ATP synthesis, this enables fine tuning of ATP synthesis as well as modulation of reactive oxygen species (ROS) and calcium ion levels (e.g. [1]). Resultantly, AOX is regarded to provide metabolic plasticity being useful for adaptation to variable biotic and abiotic stress factors (e.g. [3, 7]). Moreover, it is suggested that AOX can bypass blockade or deficiency of the MRC complexes III and IV by restoring electron flow upstream of the MRC complex III. Consequently, AOX is considered to be an important element of a therapeutic strategy against impairment of these complexes [1, 3, 9-12].

**Figure 1.**
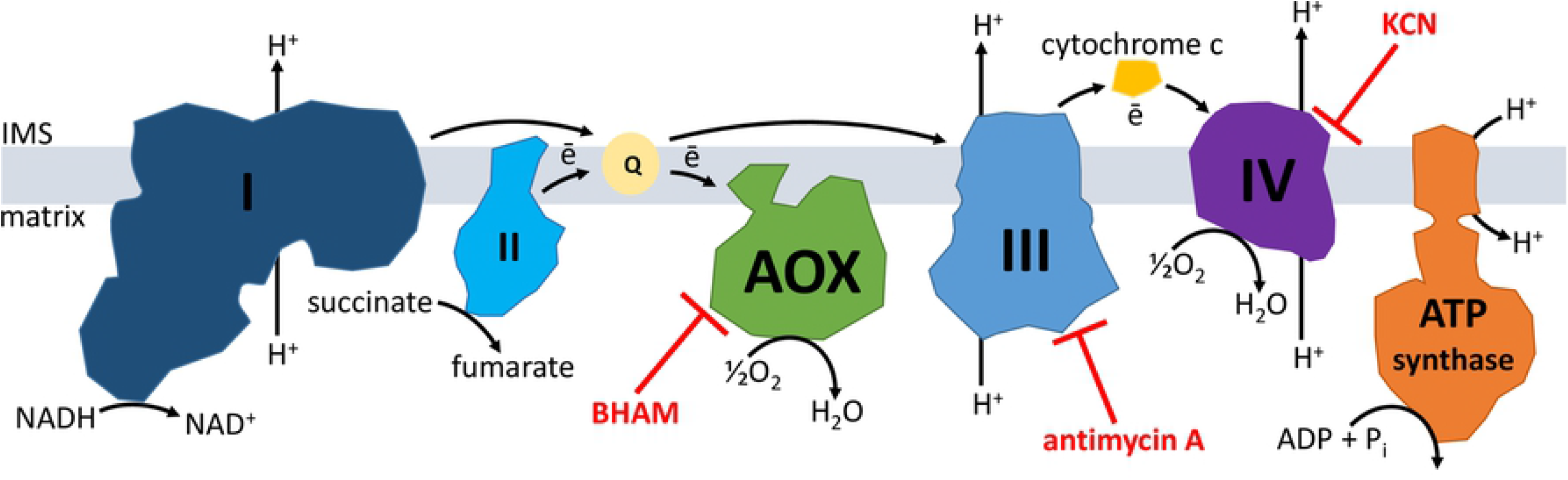
Schematic representation of the respiratory chain in animal mitochondria. The mitochondrial respiratory chain (MRC) in the inner mitochondrial membrane is formed by four main multi-subunit complexes numbered from I to IV. The presence of mitochondrial alternative oxidase (AOX) is postulated for invertebrates with exception of insects and lancelets. Electrons released due to oxidation of metabolites are transported by the MRC to oxygen and this generates proton gradient due to proton pumping by the MRC complexes I, III and IV from the matrix to the intermembrane space (IMS). The proton gradient is then used among others to feed ATP synthesis by the ATP synthase. AOX introduces a branch into the MRC localized at the ubiquinone/ubiquinol (Q) pool. This results in transferring of electrons to oxygen with sustained proton pumping by the MRC complex I, but without proton pumping by the MRC complexes III and IV. BHAM, antimycin A and KCN are known inhibitors of AOX, the MRC complex III and IV, respectively.

At present genomes of about 160 animal species representing 16 phyla, from Placozoa to Chordata with exception of insects, lancelets and vertebrates, are proposed to contain AOX encoding gene [1, 12-14]. However, the enzyme functionality was only tested in the case of a few species. The best known examples are AOXs of the pacific oyster *Crassostrea gigas* [15] and the tunicate *Ciona intestinalis* [16]. The first one was studied in isolated *C. gigas* mitochondria [17] and after heterologous expression in the yeast *Saccharomyces cerevisiae* [18] cells [19]. The second one was heterologously expressed in mitochondria of cultured human cells [9, 20-21] as well as in the fruit fly *Drosophila (Sophophora) melanogaster* [22] [23-27] and mouse [9, 12, 28-29]. Additionally, some experimental data concerning AOX protein are available for crustaceans, the brine shrimp *Artemia franciscana* [30], the white shrimp *Litopenaeus vannamei* [31] [32] and the marine copepod *Tigriopus californicus* [33] [34]. In the case of both the shrimps the data concerns AOX activity in isolated mitochondria whereas in the case of the copepod amounts of AOX mRNA and protein in various life stages of the animal and under stress temperature conditions were studied.

As it has been summarized by Rajendran et al. [9], *C. intestinalis* AOX expressed in human cells, fruit flies and mice is not active when the MRC complex III and/or IV work properly, but inhibition or overload of these complexes triggers the enzyme activity. In the case of human cell lines, *C. intestinalis* AOX expression was shown to confer spectacular cyanide resistance of mitochondrial respiration and compensate for both the growth defect and the pronounced oxidant-sensitivity caused by the MRC complex IV deficiency [20-21]. The possibility to provide a complete or substantial protection against a range of phenotypes induced by the MRC complex IV deficiency or inhibition was also observed after expression of *C. intestinalis* AOX in mitochondria of fruit flies [23-25] and mice [28-29]. The same applies to the mouse MRC complex III deficiency [9, 11]..

Thus, the available data indicates that functional studies of animal AOX are based on models that do not contain native AOX with exception of *T. californicus, C. gigas, A. franciscana* and *L. vannamei* enzyme. The molecular weight (MW) detected for the AOX protein reported for *T. californicus* is not convincing and its predicted amino acid sequence does not contain the proper C-terminus motif [34]. In the case of *C. gigas*, AOX contribution to the oyster adjustments to short-term hypoxia and re-oxygenation was considered in isolated mitochondria [17] but the presented data do not allow for clear conclusions as the observed AA-resistant respiration was not shown to be eliminated by AOX inhibitor, for example by benzohydroxamic acid (BHAM) [35]. Moreover, to observe the enzyme activity in the shrimp isolated mitochondria the authors applied an uncoupler to increase the recorded oxygen uptake rate [32]. Resultantly, the measurements were performed for uncoupled mitochondria, i.e. in the absence of ubiquinone reduction to ubiquinol, regarded as crucial for AOX activity [9, 12, 24, 29]. Thus, data concerning animal AOX expression and activity regulation is still scarce (e.g. [1, 3, 34]).

In accordance with recent sequencing data, putative AOX encoding genes have been also identified in tardigrades [1, 36]. This concerns three species, namely *Hypsibius exemplaris* [37] (former *Hypsibius dujardini* [38]), *Ramazzottius varieornatus* [39] and *Milensium inceptum* [40] (former *Milnesium tardigradum* [38]. Tardigrades are microscopic invertebrates with a body length ranging from 50 μm to 1200 μm although very few species exceed 800 μm. They inhabit marine, freshwater and terrestrial habitats, but all are regarded to be aquatic animals because they require a water-film surrounding the body to be active. The most known feature of tardigrades is the ability to enter cryptobiosis when environmental conditions are unfavorable (for review, see, for example, [40-46]).

Here, we set out to test whether AOX activity of *H. exemplaris*, as a part of the mitochondrial respiratory system, can be observed at the level of intact organisms. We also estimated whether the activity requires the MRC complex III and/or IV blockage. To that end, we determined the animal viability and respiration in the presence or absence of inhibitors of the MRC complexes and AOX as well as verified the AOX protein presence. The obtained data points at an emergence of a whole-animal model suitable to study activity and expression regulation of natively expressed animal AOX. According to our best knowledge we demonstrated, for the first time, that AOX activity of small aquatic invertebrates can be monitored by measurement of whole-body respiration due to registration of the oxygen uptake rate by intact specimen mitochondria. Moreover, we demonstrated that the enzyme contributed to animal functioning also in the absence of the MRC complex III and/or IV blockage previously described as precondition to observe animal AOX activity.

## Results

### AOX inhibitor affects H. exemplaris activity and eliminates the animal tolerance to the MRC complex III and IV inhibitors

To check functionality of the hypothetical *H. exemplaris* AOX protein (stored in the GenBank under accession number OWA52662.1), we performed *in vivo* toxicity test. We assumed that if *H. exemplaris* specimens had functional AOX, they would tolerate the presence of KCN and/or AA as well as BHAM added separately but would be affected by the combination of KCN + BHAM and AA +/- KCN + BHAM. The graphical summary of the test is shown in Fig.2A whereas Fig. 2B presents images illustrating the treated animal appearance. Treatment of *H. exemplaris* specimens with methanol (used as a solvent for BHAM and AA) for 2 h did not affect their full activity defined as coordinated movements of the body and legs as well as did not change the animal shape. In the presence of 3 mM BHAM the specimens initially did not change their activity and shape but over time their activity was moderately reduced to frequent leg movements and the animals adopted a croissant shape. The delayed changes triggered by the presence of BHAM were not reversed within 2 h of observation (or even after longer observation lasted 6 h; not shown). Addition of 1 mM KCN and 280 µg/ml AA, separately or in a sequence, almost instantaneously and significantly reduced the animal mobility that was confined to single leg movements but the animal were not paralyzed. Moreover, the shape of the animals changed into the characteristic croissant ones. Nevertheless, the animal activity was partially restored over time within 2 h of observation. In the presence of KCN and BHAM added in either order, the mobility of *H. exemplaris* specimens was completely stopped within 1 minute and the animals took the characteristic stretched/inflated shape. Furthermore, their return to activity was not detected within 2 h of observation. The rapid disappearance of full activity was also observed when BHAM was added in the presence of AA applied in a sequence with KCN. Representative images and films illustrating the animal appearance and behavior in different parts of the toxicity test are presented in Additional files 1 and 2. The obtained results suggested functionality of AOX in *H. exemplaris* mitochondria providing the animal tolerance to the MRC complex III and complex IV inhibitors. These inhibitors appeared to enhance the AOX activity in a time-dependent way. On the other hand, the complex blockage was not indispensable for observation of AOX contribution to animal functioning.

**Figure 2.**
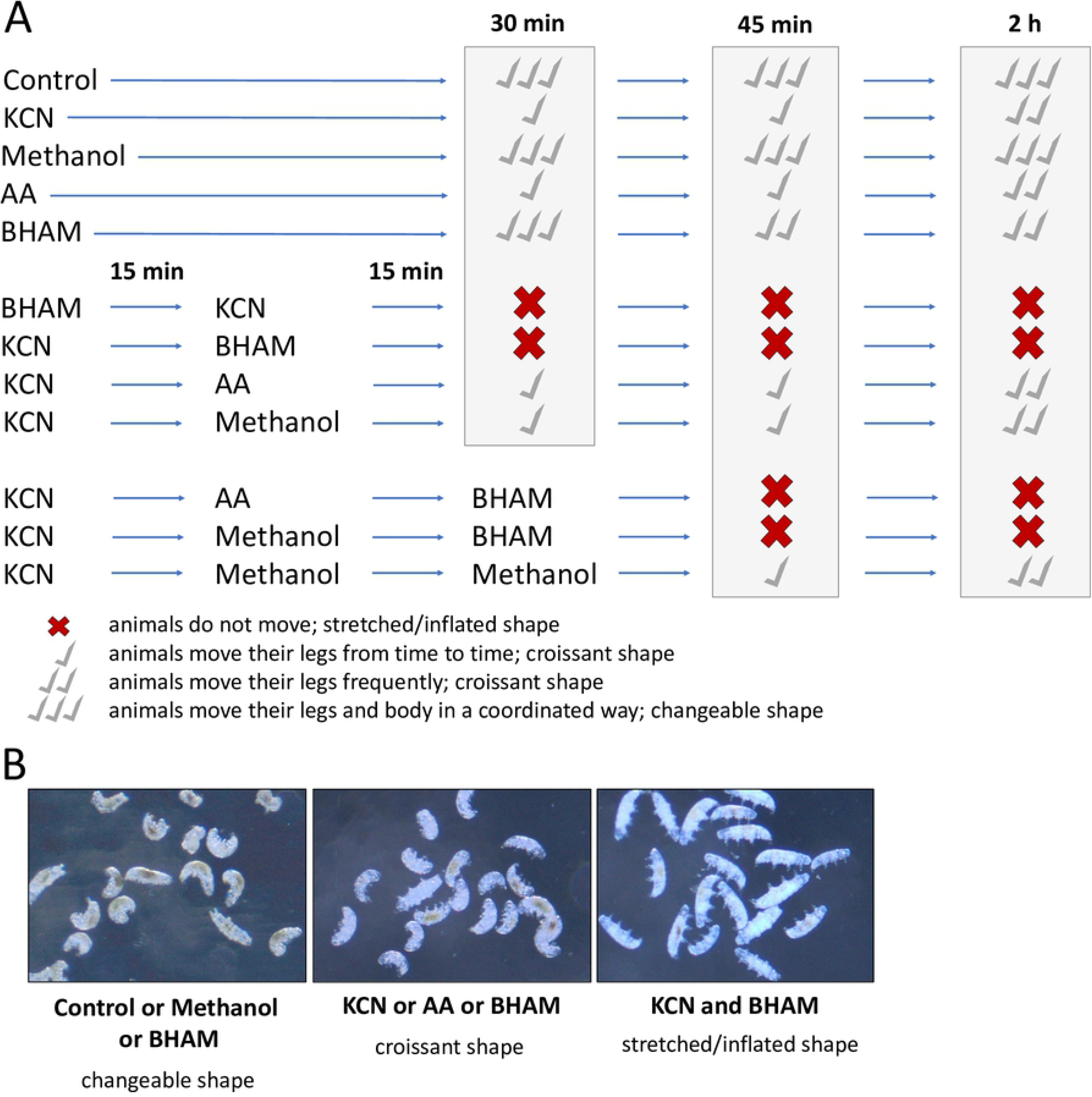
Toxicity in vivo test indicates AOX contribution to functioning of *H. exemplaris* specimens in the absence and in the presence of the MRC complex III and IV inhibitors (AA and KCN, respectively). Adult active specimens of a comparable body length and cleaned of debris were treated with AA, KCN and BHAM in different configurations and observed for 2h in the indicated time points. The applied concentrations of inhibitors were as follows 280 µg/ml AA, 3 mM BHAM and 1 mM KCN. Methanol was used as a solvent control for AA and BHAM. (A) Graphic representation of the treated animal activity. (B) The treated animal body shape changes co-occurring with mobility changes. The data represent two independent experiments.

### AOX inbibitor-sensitive respiration enables by-pass of the MRC complex III and IV inhibition in H. exemplaris mitochondria but can be also observed in the absence of the inhibition

To verify AOX activity contribution to *H. exemplaris* tolerance to the MRC complex III and IV inhibitors, we measured the rate of oxygen uptake by mitochondria of intact specimens under the conditions corresponding to the *in vivo* toxicity test. As shown in Fig. 3A, mitochondria-based respiration of intact specimens was easily measured and affected by the inhibitors of MRC and/or AOX (see also Additional file 3 for the raw data). The combination of KCN with BHAM caused the respiration elimination independently of the presence of AA (Fig. 3B). Addition of KCN +/- AA resulted in the respiration inhibition close to 60% of the initial respiration and the remaining respiration was sensitive to AOX inhibitor (BHAM). In the absence of KCN, the recorded respiration was slightly sensitive to BHAM and the effect of the inhibitor was delayed. Thus, the BHAM-sensitive oxygen uptake by *H. exemplaris* intact specimen mitochondria could be observed without the blockage of the MRC complexes III and/or IV but the blockage was required for the respiration to be pronounced. Accordingly, mass spectrometry allowed for detection of AOX protein only in the protein extract prepared from specimens treated for 2 h with KCN (see also see also Additional file 4 for the detection data) that implies a role of the MRC cytochrome pathway in modulation of *H. exemplaris* AOX amount. It should be also mentioned that the AOX protein was detected in MW range of 35-40 kDa which corresponds to heterologously expressed animal AOX proteins [19].

**Figure 3.**
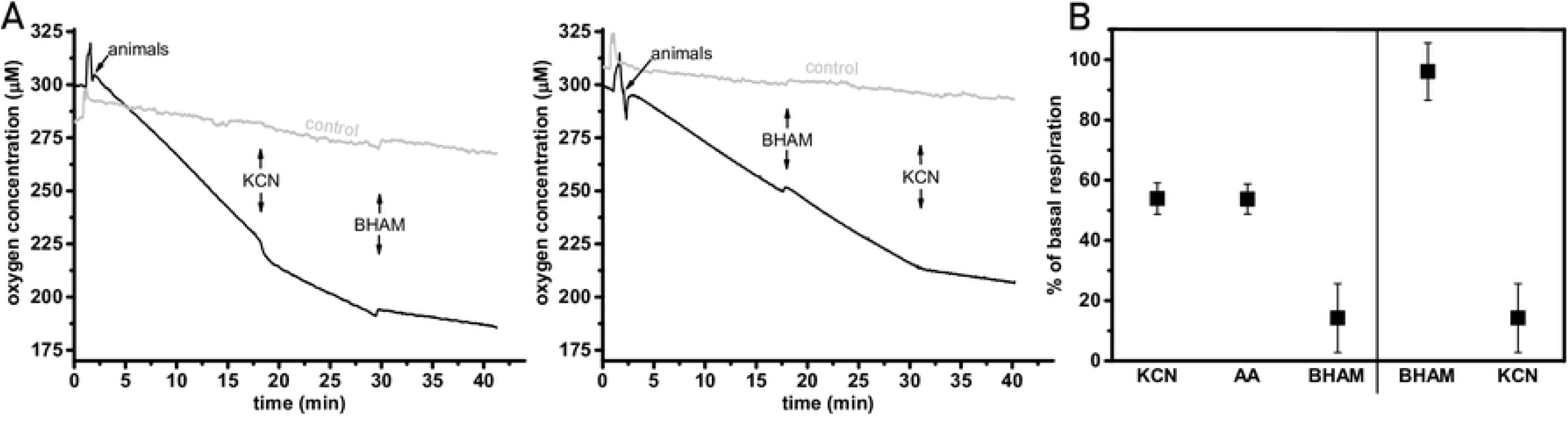
The measurements of the rate of oxygen uptake by intact *H. exemplaris* specimens confirm the tardigrade AOX activity. (A) Representative traces of the performed measurements of the rate of oxygen uptake. Control denotes the measurements made for the proper volume of the culture medium taken from the animal suspension. (B) Changes in *H. exemplaris* specimen respiration in the presence of inhibitors of AOX (BHAM) as well as the MRC complex III (AA) and complex IV (KCN). The applied concentrations of inhibitors were as follows 280 µg/ml AA, 3 mM BHAM and 1 mM KCN. The data are presented as mean values ± SD.

## Discussion

Here, we report that the tardigrade *H. exemplaris* possesses functional AOX that contributes to specimen tolerance to the mitochondrial respiratory chain (MRC) complex III and IV inhibition. Simultaneously, according to our best knowledge, it is the first report showing that small aquatic invertebrate AOX activity can be monitored by measurement of the oxygen uptake rate by mitochondria of intact specimens reflecting whole-body respiration. This makes the tardigrade a promising *in vivo* model that is amenable to whole-organism analyses. So far two other animal models, both obtained due to heterologous expression of the tunicate *C. intestinalis* AOX in fruit flies and mice, have been used in research on AOX impact on animal physiology and the enzyme possible application in therapy of human mitochondrial diseases (e.g. [1, 3, 9-12]. The data obtained for *H. exemplaris* could supplement the models due to the presence of native AOX as well as simplicity of the tardigrade culture and AOX research in intact specimens.

The common property of *C. intestinalis* AOX heterologously expressed in fruit fly and mouse mitochondria is that it confers a substantial tolerance to MRC inhibitors *in vivo* [23, 28-29] but becomes enzymatically active only when MRC becomes inhibited beyond ubiquinone [9, 20-21, 24]. In the case of natively expressed *H. exemplaris* AOX the specimen BHAM-sensitive respiration is observed in the presence of previously added KCN and/or AA. Accordingly, *H. exemplaris* specimens treated with the MRC complex III and/or IV inhibitors after initial restrained mobility partially return to activity within 2 h and this return is prevented by the presence of BHAM. The results of the performed *in vivo* toxicity test are corroborated by the possibility of AOX protein detection after *H. exemplaris* specimen treatment with KCN for 2h. Moreover the results suggest that *H. exemplaris* specimens are probably more tolerant to KCN treatment than AOX-expressing fruit flies. The latter remained active during 20 - 30 min of the KCN treatment and then recovered from paralysis overnight [23]. It has also been shown that the heterologous AOX expression in mice enables the anesthetized animal survival after exposition to a lethal concentration of gaseous cyanide [28] or injected KCN [29]. It is clear that without the use of anesthesia, these experiments would have been inhumane although it should be remembered that anesthetics may contribute to mitochondrial dysfunction [28, 47].

The blockage of MRC at the level of the complexes III and/or IV results in hypoxia or even anoxia described as important factors controlling the animal AOX transcript levels [3, 17, 48]. On the other hand, AOX affinity to its quinone substrate may increase under condition of the substrate prolonged reduction resulting from the blockage of the MRC complexes III and/or IV (e.g. [27]). However, in the case of *H. exemplaris*, the specimen activity and respiration are sensitive to BHAM also in the absence of MRC complex III and/or IV inhibition. Namely, after the BHAM treatment, slight and delayed inhibition of the oxygen uptake is observed that corroborates to the delayed change in the treated animal shape and moderate mobility restriction. This allows for assumption that mechanism of the animal AOX activity regulation is more complex. Conversely, the amount of AOX in *H. exemplaris* specimens increases distinctly, above the threshold of detection by mass spectrometry, when these animals appear to cope with the presence of KCN. This in turn indicates at important mechanism of the encoding gene regulation at protein level imposed by the blockage of the MRC complex IV that cannot be fully studied using animals with the AOX heterologous expression controlled in an artificial way.

## Conclusions

As has been mentioned by Saari et al. [3], despite the encouraging findings to date in animal models, the potential problems in the use of AOX in human disease therapy need to be considered in much greater detail. Accordingly, application of *H. exemplaris*, could help to explain inherent properties of natively expressed AOX, in particular the encoding gene expression and the enzyme activation, and how the activation affects the organism. Thus, we propose a whole-organism animal model for research on natively expressed AOX that enables analysis of the enzyme contribution to animal behavior and mitochondria functionality under both physiological conditions and a given biochemical defect or external stress. It is also important that protocols for *H. exemplaris* gene expression and genetic manipulation were developed (e.g. [49-50]) that increases the range of possible experiments.

## Materials and Methods

### Taxonomy

Species citation follows International Code of Zoological Nomenclature. Tardigrade taxonomy follows Bertolani et al. [51] and later updates for Isohypsibiidae [52].

### Reagents

The following inhibitors were applied: benzohydroxamic acid (BHAM; #412260) for AOX, potassium cyanide (KCN; #60178) for the MRC complex IV and antimycin A (AA; #A8674) for the MRC complex III (purchased from Sigma-Aldrich).

### Culture of Hypsibius exemplaris

*Hypsibius exemplaris* Z151 strain was purchased from Sciento (Manchester, United Kingdom). To maintain the culture, specimens were kept in POL EKO KK 115 TOP^+^ climate chamber (photoperiod 12h light /12h dark, 18 °C, relative humidity of 50%) on Petri-dishes (55 mm in diameter) with bottom scratched by sandpaper to allow movement of tardigrades. They were covered with a thin layer of the culture medium obtained by mixing double-distilled water and spring water (Zywiec Zdroj S.A., Poland) in ratio of 3 to 1. *Chlorella vulgaris* SAG211-11b strain was served as a food once per week after the plate cleaning. Animals were transferred to new culture plates every few months. The SAG211-11b strain was a kind gift of Marcin Dziuba (Department of Hydrobiology, Faculty of Biology, Adam Mickiewicz University, Poznań, Poland) and was obtained from the culture collection of algae (Sammlung von Algenkulturen (SAG)) at the University of Göttingen, Germany.

### Toxicity in vivo test

The test was performed in glass blocks with hemispherical cavity of 32 mm in diameter (Karl Hecht Staining Blocks Molded Glass 2020, Lab Unlimited). Only adult and fully active (displaying coordinated movements of the body and legs) specimens with a body length of about 200 µm and cleaned of debris were collected 16 h before the test beginning and kept in the glass block in 3 ml of the culture medium. To check the impact of KCN, AA and BHAM, groups of 20 fully active specimens were transferred into 1 ml of the culture medium in new glass blocks. Then they were treated with combinations of KCN, BHAM, AA and methanol (used as a solvent for AA and BHAM) for 2 h at 18 °C and observed in a few time points (i.e. 30 min, 45 min and 2 h). The applied concentrations of these inhibitors were established experimentally to obtain saturated effect on the oxygen uptake rate inhibition (see Measurements of *H. exemplaris* respiration) and were as follows: 1 mM KCN, 3 mM BHAM and 280 µg/ml AA. The test was repeated two times. The specimens activity before and after the treatment was monitored under Olympus SZ61 stereo microscope connected to Olympus UC30 microscope digital camera. Images and short video films were obtained using Olympus CellSens Standard Software.

### Measurements of H. exemplaris respiration

To estimate animal respiration, the rate of oxygen uptake was measured at 18 °C in 0.5 ml of the culture medium, using two water-thermostated incubation chambers with computer-controlled Clark-type O_2_ electrode (Oxygraph, Hansatech, UK). The adult fully active specimens (about 200 µm in a body length) were collected as described above for Toxicity *in vivo* test in the amount of maximally 2000 specimens per each glass block. To obtain specimens with empty gut, (to avoid an impact of AOX belonging to *C. vulgaris* applied as a food) they were starved for three days and the culture medium was exchanged once per day. Then, the specimens were transferred to a glass test-tube, and kept in 10 ml of the culture medium. The medium was replaced 3-4 times before the measurements to avoid hypoxia. A trace was recorded for a suspension of 10 000 *H. exemplaris* specimens (without algae in the gut) in 0.1 ml of the glass test-tube medium. The proper control trace was performed simultaneously in the second chamber in the absence of animals, but with addition of 0.1 ml of the glass test-tube medium and the applied reagents. To avoid hypoxia, the oxygen uptake rate measurements were finished when oxygen level achieved 50% of its maximal value. The measurements were performed at least in triplicate. The statistical significance of results was tested using unpaired t-test at the 0.05 level of significance.

### Detection of AOX protein

The presence of AOX protein was estimated by liquid chromatography coupled to tandem mass spectrometry (LC–MS/MS) performed in the Department of Biomedical Proteomics, Institute of Bioorganic Chemistry, Polish Academy of Sciences (Poznań, Poland). Amino acid sequences annotated as animal AOX and stored in the GenBank were used for the presence detection. The analyzed samples were cut from SDS-PAGE gel at its part corresponding to MW between 25-34 and 35-40 kDa. Each sample was obtained for 15 000 adult fully active specimens with empty gut incubated for 2 h at 18 °C in 1 ml of the culture medium in the glass cubes in the absence or in the presence of 1 mM KCN. Then the animals were pelleted in 1.5 ml Eppendorf tubes (8 000 g for 2 min), and the pellet was frozen in liquid nitrogen and stored at -80 °C until protein extraction. To obtain the extract, 50 µl of an isolation buffer (50 mM ammonium bicarbonate, 1% SDS, 1x Protein Inhibitor Thermo Fisher Scientific, #78430) were added to each tube containing the pellet. Then the samples were frozen and thawed three times, sonicated using Bioruptor Plus UCD-300 (4 °C, 20 kHz, 15 × 30 s. with 30 s interval between each sonication pulse) and centrifuged (8 000 g for 2 min in 4 °C). The obtained supernatant (app. 50 µl) was transferred from each Eppendorf tube to the new one. Total amount of proteins in each of the supernatants was estimated by Bradford assay (5 replications). 60 µg of each sample was denaturated using Sigma 2x lysis buffer (#3401) for 5 min in 95 °C and loaded on SDS-PAGE gel. Protein separation in 14% resolving gel was performed according to standard protocol [53].

## Additional files

**Additional file 1: Figure S1**. Images of animals after 30 min and 2 h of all variants of the performed toxicity *in vivo* test. File format: pdf.

**Additional file 2: Movie S1**. Recorded films presenting animals after 45 min treatment with BHAM, KCN and BHAM + KCN as well as without the treatment. File format: video.mp4.

**Additional file 3: Table S1**. Raw data used to calculate the impact of AOX and the MRC complex III and IV inhibitors on mitochondria-based respiration of intact specimens. File format: docx.

**Additional file 4: Table S2**. LC-MS/MS based detection of *H. exemplaris* AOX protein. File format: docx.

## List of abbreviations

AA: antimycin A
AOX: mitochondrial alternative oxidase
BHAM: benzohydroxamic acid
KCN: potassium cyanide
MRC: mitochondrial respiratory chain

## Ethics approval and consent to participate

Not applicable.

## Consent for publication

Not applicable.

## Availability of data and materials

Data generated and analyzed during this study are included in this published article and its supplementary information files.

## Authors’ contributions

DG, AK, LK and HK came up with research ideas. HK, WJ and LK supervised the performed analyses. DG, MR, LK, AK and HK wrote the final version of the manuscript. DG, MR and LK carried out the tardigrade cultures and collected animals for experiments. DG and MR performed toxicity *in vivo* test and oxygen uptake rate measurements. DG prepared samples for LC–MS/MS analysis as well participate with AK with result analysis. All authors read and approved the final manuscript.

## Acknowledgements

We are extremely grateful to Magdalena łuczak (Department of Biomedical Proteomics, Institute of Bioorganic Chemistry, Polish Academy of Sciences, Poznan, Poland) for her invaluable contribution to LC–MS/MS analysis. We would also like to thank to Sławek Cerbin and Marcin K. Dziuba for *Chlorella vulgaris* SAG211-11b strain used to feed the cultured *H. exemplaris* specimens.

